# Comparing protein-protein interaction networks of SARS-CoV-2 and (H1N1) influenza using topological features

**DOI:** 10.1101/2021.11.02.463717

**Authors:** Hakimeh Khojasteh, Alireza Khanteymoori, Mohammad Hossein Olyaee

## Abstract

**Background:** SARS-CoV-2 pandemic first emerged in late 2019 in China. It has since infected more than 183 million individuals and caused about 4 million deaths globally. A protein-protein interaction network (PPIN) and its analysis can provide insight into the behavior of cells and lead to advance the procedure of drug discovery. The identification of essential proteins is crucial to understand for cellular survival. There are many centrality measures to detect influential nodes in complex networks. Since SARS-CoV-2 and (H1N1) influenza PPINs pose 553 common proteins. Analyzing influential proteins and comparing these networks together can be an effective step helping biologists in drug design.

**Results:** We used 21 centrality measures on SARS-CoV-2 and (H1N1) influenza PPINs to identify essential proteins. PCA-based dimensionality reduction was applied on normalized centrality values. Some measures demonstrated a high level of contribution in comparison to others in both PPINs, like Barycenter, Decay, Diffusion degree, Closeness (Freeman), Closeness (Latora), Lin, Radiality, and Residual. Using validation measures, the appropriate clustering method was chosen for centrality measures. We also investigated some graph theory-based properties like the power law, exponential distribution, and robustness.

**Conclusions:** Through analysis and comparison, both networks exhibited remarkable experimental results. The network diameters were equal and in terms of heterogeneity, SARS-CoV-2 PPIN tends to be more heterogeneous. Both networks under study display a typical power-law degree distribution. Dimensionality reduction and unsupervised learning methods were so effective to reveal appropriate centrality measures.

## Background

SARS-CoV-2, a novel coronavirus mostly known as Covid-19, has become a matter of critical concern for every country around the world. It was first identified in December 2019 in Wuhan, China. The coronavirus Covid-19 is affecting 220 countries and territories around the world. As of 2 July 2021, over 183 million cases have been confirmed, with about 4 million confirmed deaths attributed to COVID-19.

In the past several years, extensive experiments and data evolution have provided a good opportunity for systematic analysis and a comprehensive understanding of the topology of biological networks and biochemical processes in the cell [1]. In other words, we need to choose the right essential proteins to be attacked by the new drug. However, identifying appropriate drug target proteins through experimental methods is time-consuming and expensive [2, 3]. Both SARS-CoV-2 and (H1N1) influenza viruses have similar clinical symptoms [4]. SARS-CoV-2 and (H1N1) influenza PPINs have 553 common proteins. Essential proteins play a vital role in the survival and development of the cell. They are also the most important materials in a variety of life processes. In cellular life, the importance of different protein activities is not the same. The identification of essential proteins is decisive to understanding the minimal requirements for cellular life and for practical purposes, such as a better understanding of diseases, and drug discovery [5]. Studying SARS-CoV-2 and (H1N1) influenza PPINs can be helpful to investigate similarities and differences between them. Studies have shown that PPIs involve more heterogeneous processes and pose a range of sizeable regulations. For a more accurate understanding of their importance in cell life, it has to identify various interactions and determine the consequences of the interactions[6]. Moreover, this can use to empirically investigate complex network properties such as degree distribution [7], power-law [8], and other topological features.

Hahn et al. [9] examined essential proteins in PPINs of eukaryotes: yeast, worm, and fly through three centrality measures. The results showed that there was a clear relationship between central proteins and survival. To detect which centrality measure is more suitable for choosing essential proteins in PPINs, Ernesto[10] investigated the relationships between several centrality measures and subgraph centrality with essential proteins in the yeast PPIN. His study indicates that protein essentiality appears to be related to how much a protein is involved in clusters of proteins. As a result, subgraph centrality outperformed better than other measures for detecting essential proteins. Ashtiani et al. surveyed 27 centrality measures on yeast protein-protein interaction networks for ranking the nodes in all PPINs. They examined the correlation between centrality measures through unsupervised machine learning methods. [11].

In this work, we adopted the human PPIN data set from [12, 13] database to compare SARS-CoV-2 and (H1N1) influenza PPINs. Using these networks, we then analyzed the topological features, focusing on the properties of the graphs which represent these networks. We considered some specific measures, such as graph density, degree distribution, and 21 different centrality measures. We fitted power law and exponential distributions on these networks and calculated alpha power and R-squared values.

## Methods

We propose a useful analysis approach to compare SARS-CoV-2 and (H1N1) influenza PPINs. At first, we need to select a valid dataset and so, investigate and select suitable features that are meaningful in a biological system. Next, we develop our approach to make comparisons and the results are analyzed. In the following, we describe how to deal with these phases, respectively. The process start by computing global network properties. In the next phase, 21 different centrality measures are applied on both networks, standard normalization and PCA are used on centrality values, respectively. Using some machine learning methods, the centrality measures are compared and analyzed.

### Network Global properties

In this study, we have considered some of the network properties such as graph density, graph diameter, and centralization. In the following, we review these network concepts. All these properties are calculated and analyzed in both networks using igraph [14] R package. Then, the power-law distribution is checked out by computing α and R-squared values. R-squared is the percentage of the response variable variation that is described by a linear model [15].

Although, PPINs are directed but most of analyzing methods consider PPINs as undirected [16]. For this research study, we considered PPINs as undirected and loop-free connected graphs. So, let *G* = (*V*, *E*) be an undirected graph. This graph consists of nodes represented by *V* = {*ν*_1_, *ν*_2_ … } and edges *E* = {*e*_1_, *e*_2_ … } such that any edge *e*_*ij*_ ∈ *E* represents the connection between nodes *ν*_*i*_ and *ν*_*j*_ ∈ *V*.

#### Graph density

The density of a graph is the fraction of the number of edges to the number of possible edges [17]. Density is equal to 2 ∗ |*E*| divided by |*V*| ∗ (|*V*| − 1). A complete graph has density 1; the minimal density of any graph is 0. There are some features for identifying biological networks. Often, biological networks are incomplete or heterogeneous which means very low density [18].

#### Graph diameter

In a network, diameter is the longest shortest path between any two vertices (*u*, *ν*), where *d*(*u*, *ν*) is a graph distance[19].

#### Heterogeneity

The network heterogeneity is defined as the coefficient of variation of the connectivity distribution:

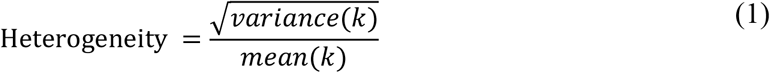

In PPINs, the connectivity *k*_*i*_ of node *i* equals the number of directly linked neighbors. PPINs tend to be very heterogeneous. Highly connected ‘hub’ nodes in PPINs have an important role in the network. A hub protein is essential and contains many distinct binding sites to accommodate non-hub proteins [20].

#### Centralization

Centralization is a method that give information about topology of a network. Centralization is measured from the centrality scores of the vertices. The centralization that closes to 1, illustrates that probably the network has a star-like topology. If it is closer to 0, the more likely topology of the network is like square whereas every node of the network has at least 2 neighbors) [19]. This metric is calculated as follows [21]:

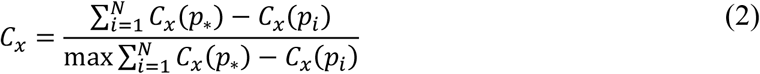

where *C*_*x*_(*p*_*i*_) is any centrality measure of point *i* and *C*_*x*_(*p*_∗_) is the largest such measure in the network. Each centrality measure can be used (betweenness centrality, closeness centrality and etc).

#### Centrality analysis

In this work, the following 21 centrality measures are selected: Average Distance [22], Barycenter [23], Closeness (Freeman) [21], Closeness (Latora) [24], Residual closeness [25], Decay [26], Diffusion degree [27], Geodesic K-Path [28, 29], Laplacian [30], leverage [31], Lin [32], Lobby [33], Markov [34], Radiality [35], Eigenvector [36], Subgraph scores [37], Shortest-Paths betweenness [21], Eccentricity [38], Degree [19], Kleinberg’s authority scores [39], and Kleinberg’s hub scores [39]. These measures are calculated using the centiserve [40] and igraph [14] R packages. We have classified the centrality measures into five distinct classes including Distance-, Degree-, Eigen-, Neighborhood-based and miscellaneous groups depend on their logic and formulas (Table 1).

**Table 1.** Centrality measures. The centrality measures are classified in five groups depending on their logic and formula.

| Distance based | Degree based | Eigen based | Neighborhood based | Miscellaneous |
| --- | --- | --- | --- | --- |
| Average Distance | Kleinberg's authority centrality scores | Eigenvector Centrality Scores | Subgraph centrality scores | Geodesic K-Path Centrality |
| Barycenter | Degree Centrality | Laplacian Centrality |  | Markov Centrality |
| Closeness Centrality (Freeman) | Diffusion Degree |  |  | Shortest-Paths Betweenness Centrality |
| Closeness Centrality (Latora) | Kleinberg's hub centrality scores |  |  |  |
| Decay Centrality | Leverage Centrality |  |  |  |
| Eccentricity | Lobby Index (Centrality) |  |  |  |
| Lin Centrality |  |  |  |  |
| Radiality Centrality |  |  |  |  |
| Residual Closeness centrality |  |  |  |  |

Tables 2 and 3 show the definitions for 21 different centrality measures based on their group.

**Table 2:**
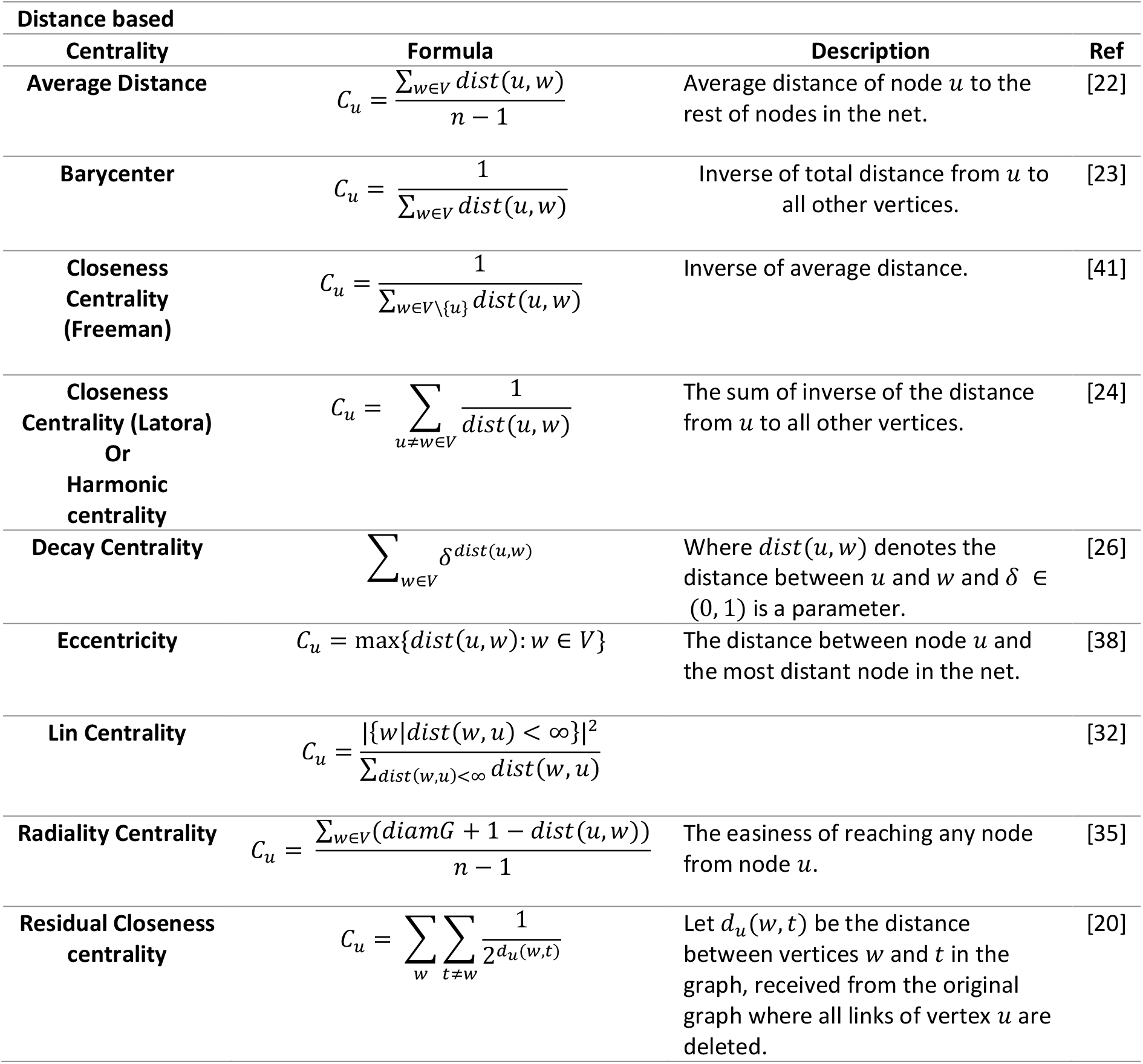
Definitions for distance based centrality measures.

**Table 3:**
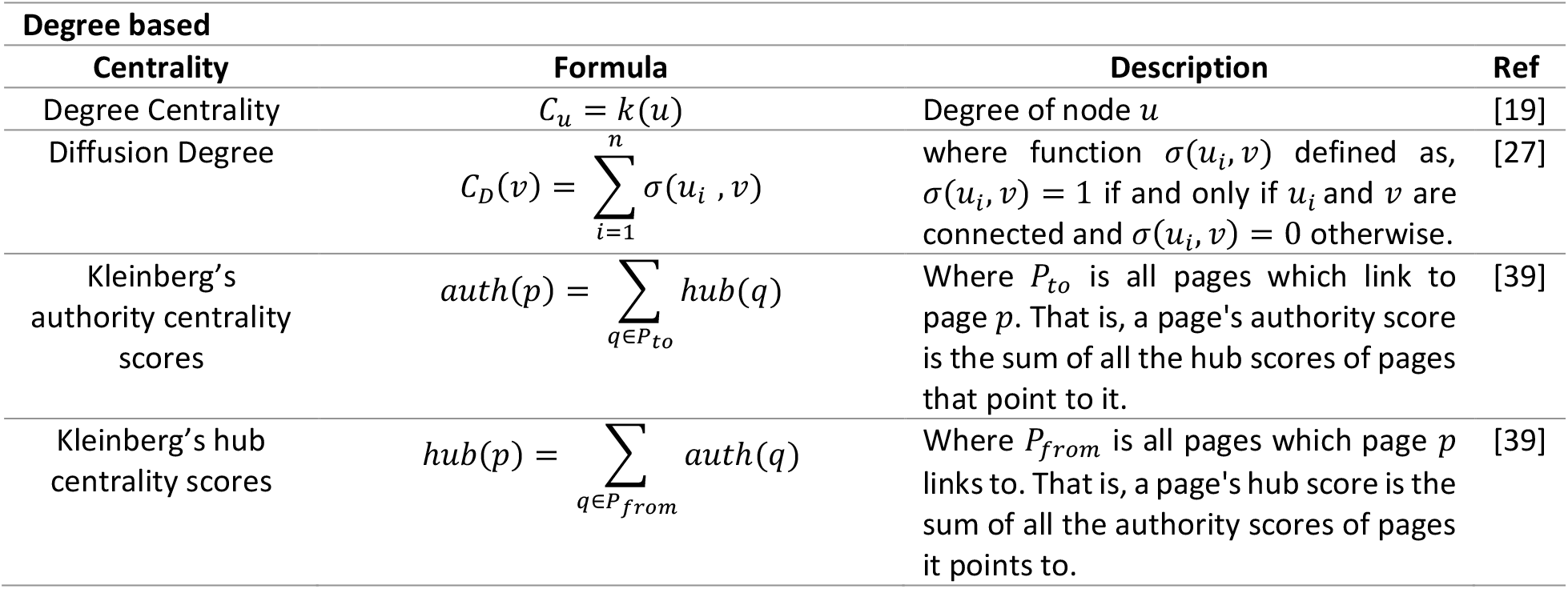
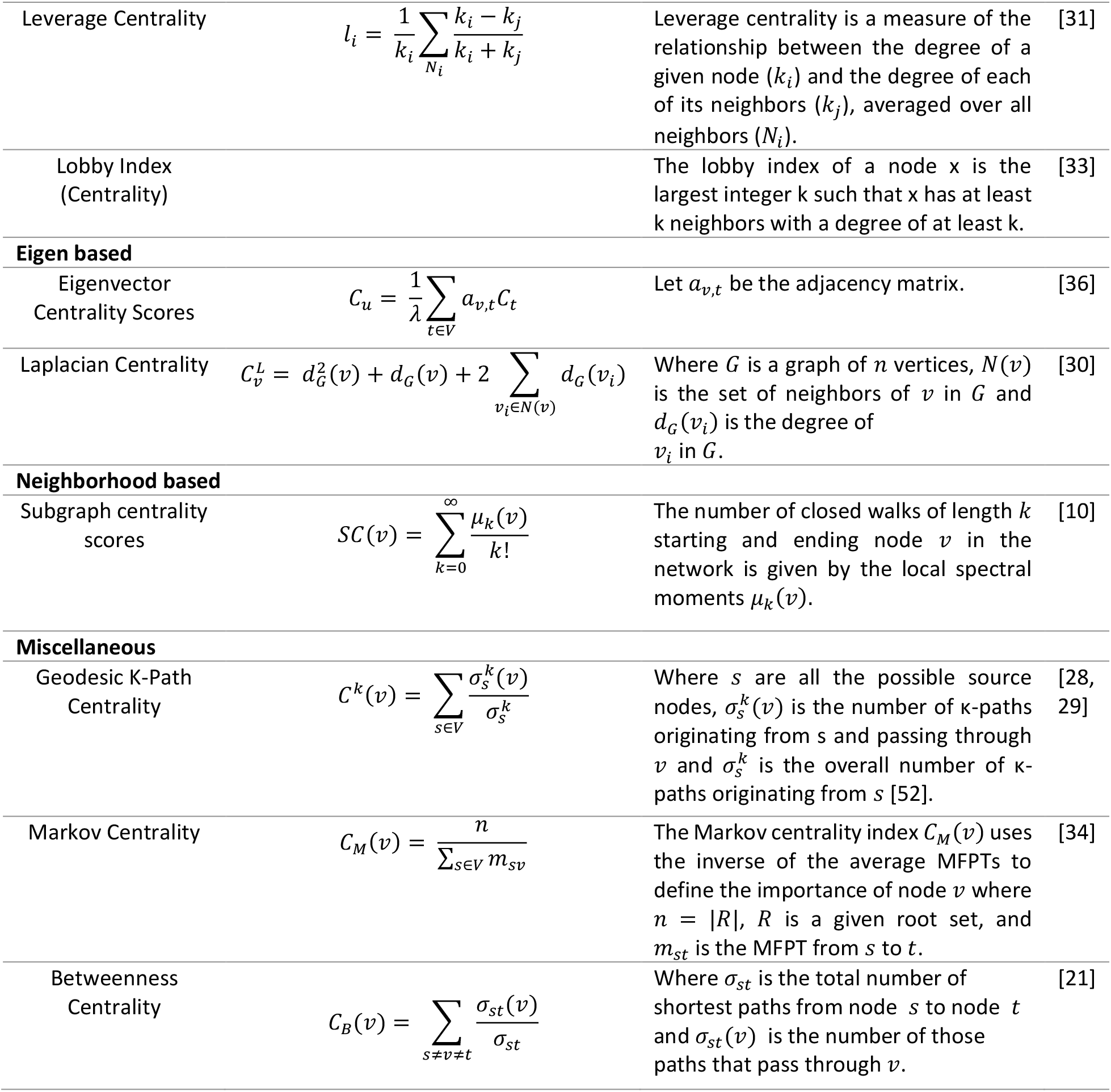
Definitions for degree based, eigen based, neighborhood based, and miscellaneous centrality measures.

### Unsupervised machine learning analysis

PCA (Principal Component Analysis) is a dimensionality-reduction method that is often used to reduce the dimensionality of large data sets, by linear transforming a large set of variables into smaller ones [42]. The aim of PCA is to remove correlated centralities, reduce overfitting, and better visualization. Since the values of centrality measures are in different scales and PCA is affected by scale, Standard normalization has been undertaken on centrality measures before applying PCA. This phase is significant because it helps to recognize which centrality measures can determine influence nodes within a network. Then, PCA is used on normalized computed centrality measures. In the next phase, it is assessed that whether it is feasible to cluster the centrality measures in both networks according to clustering tendency. Before applying any clustering method on the dataset, it’s important to evaluate whether the data sets contain meaningful clusters or not. For assessment of the feasibility of the clustering analysis, the Hopkins’ statistic values and visualizing VAT (Visual Assessment of cluster Tendency) plots are calculated by factoextra R package [43]. Some validation measures are used to select the most suitable clustering method among hierarchical, k-means, and PAM (Partitioning Around Medoids) methods using clValid package [44]. In this study, we apply Silhouette scores to select the appropriate method. After the choice of the clustering method, factoextra package is employed to find the optimal number of clusters [43]. In the clustering procedure, Wards Method [45] is used as a dissimilarity measure. Ward’s minimum variance method creates groups such that variance is minimized within clusters.

### Datasets

There are four different types of Coronaviruses (CoVs) includes Alphacoronoavirus, Betacoronavirus, Deltacoronavirus, and Gammacoronavirus [12]. Betacoronavirus includes five subtypes among Embecovirus, Sarbecovirus, Merbecovirus, Nobecovirus, and Hibecovirus. SARS-CoV and SARS-CoV-2 are from Sarbecovirus (SV) subgenus. Khorsand et al. [12] created Sarbecovirus-human protein-protein interaction network. We have derived SARS-CoV-2 PPINs from this dataset. For (H1N1) influenza PPIN, Khorsand et al. [13] made Comprehensive PPINs for all genres of Alphainfluenza viruses (IAV). The main human influenza pathogens are Alphainfluenza viruses (IAV) that include subtypes of combining one of the 16 hemagglutinin (HA: H1– H16) with one of the 9 neuraminidase (NA: N1–N9) surface antigens. We have downloaded the whole network and separated (H1N1) influenza PPIN from the Alphainfluenza protein–protein interaction network.

## Results

### Evaluation of network properties

In this study, both networks were examined to compare global properties. The network global properties were computed for both networks (Table 4). The densities of SARS-CoV-2 and (H1N1) influenza PPINs were computed at 0.0019 and 0.0023 that was expected because biological networks are usually sparse. The network diameters were equal in both networks. SARS-CoV-2 and (H1N1) influenza PPINs were correlated to the power-law distribution with high alpha power and R-squared values. In terms of comparison of heterogeneity values, SARS-CoV-2 PPIN achieved a higher value. But, both networks are relatively heterogeneous. Values of network centralization were very close together. Fig. 1. demonstrates power law (red curve) and exponential (blue curve) distributions in SARS-CoV-2 and (H1N1) influenza PPINs. Both the degree distributions were left-skewed analogous to scale-free networks.

**Table 4.** Network global properties of SARS-CoV-2 and (H1N1) influenza PPINs.

| Networks Properties | Nodes | Edges | Density | Diameter | $\alpha$ value (Power Law) | R-squared (Power Law) | Heterogeneity | Network Centralization |
| --- | --- | --- | --- | --- | --- | --- | --- | --- |
| SARS-CoV-2 | 1409 | 1922 | 0.0019 | 6 | 0.805 | 0.54 | 7.3628 | 0.3089 |
| (H1N1) influenza | 2797 | 9174 | 0.0023 | 6 | 1.009 | 0.717 | 5.3197 | 0.2392 |

**Fig. 1.**
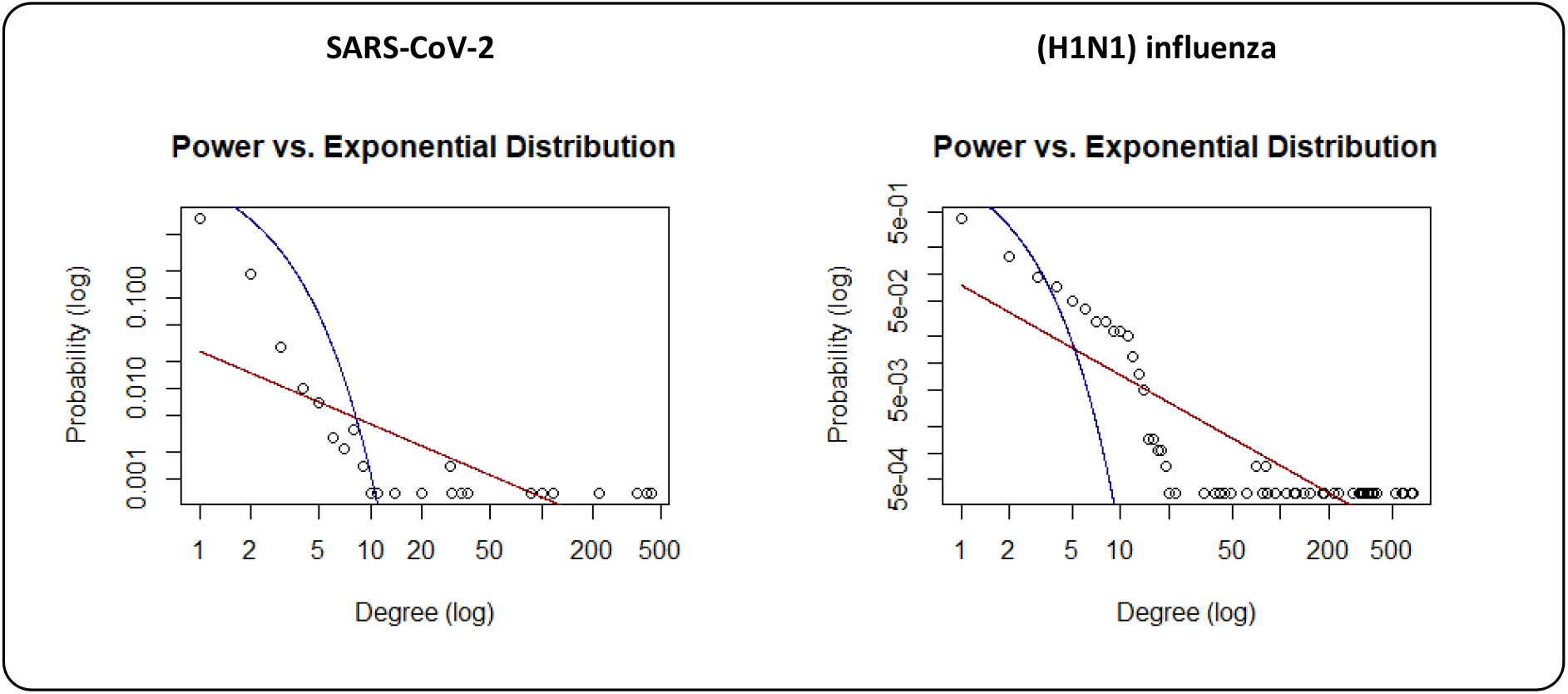
Fitting both SARS-CoV-2 and (H1N1) influenza PPINs on power-law distribution.

### Centrality analysis

In the next phase, the 21 centrality measures of nodes were calculated in the both networks. The centrality measures were divided into two groups according to Table 2: (1) distance based and (2) degree based, eigen based, and neighborhood based. R Pearson correlation measures a linear dependence between two variables (a and b). The r Pearson correlation coefficients between centralities in two groups and pairwise scatter plots of centrality measures were also shown in Fig. 2. These plots illustrate that there is a clear correlation in some of the centrality measures. For a better comparison, we also provided the dissimilarity matrix based on the Pearson correlation coefficient for all centrality measures in both networks (Fig. 3.) The range of Pearson correlations is between 0 and 2. A value of 0 indicates that there is no association between the two centrality measures. In both networks, the matrixes indicate a high positive association between Average Distance and Radiality centrality measures are highly associated together. Furthermore, in (H1N1) influenza, these correlations are more obvious between Average Distance and Lin, Barycenter, Closeness (Freeman), Radiality, Closeness (Latora), Residual closeness, and Decay measures.

**Fig.2.**
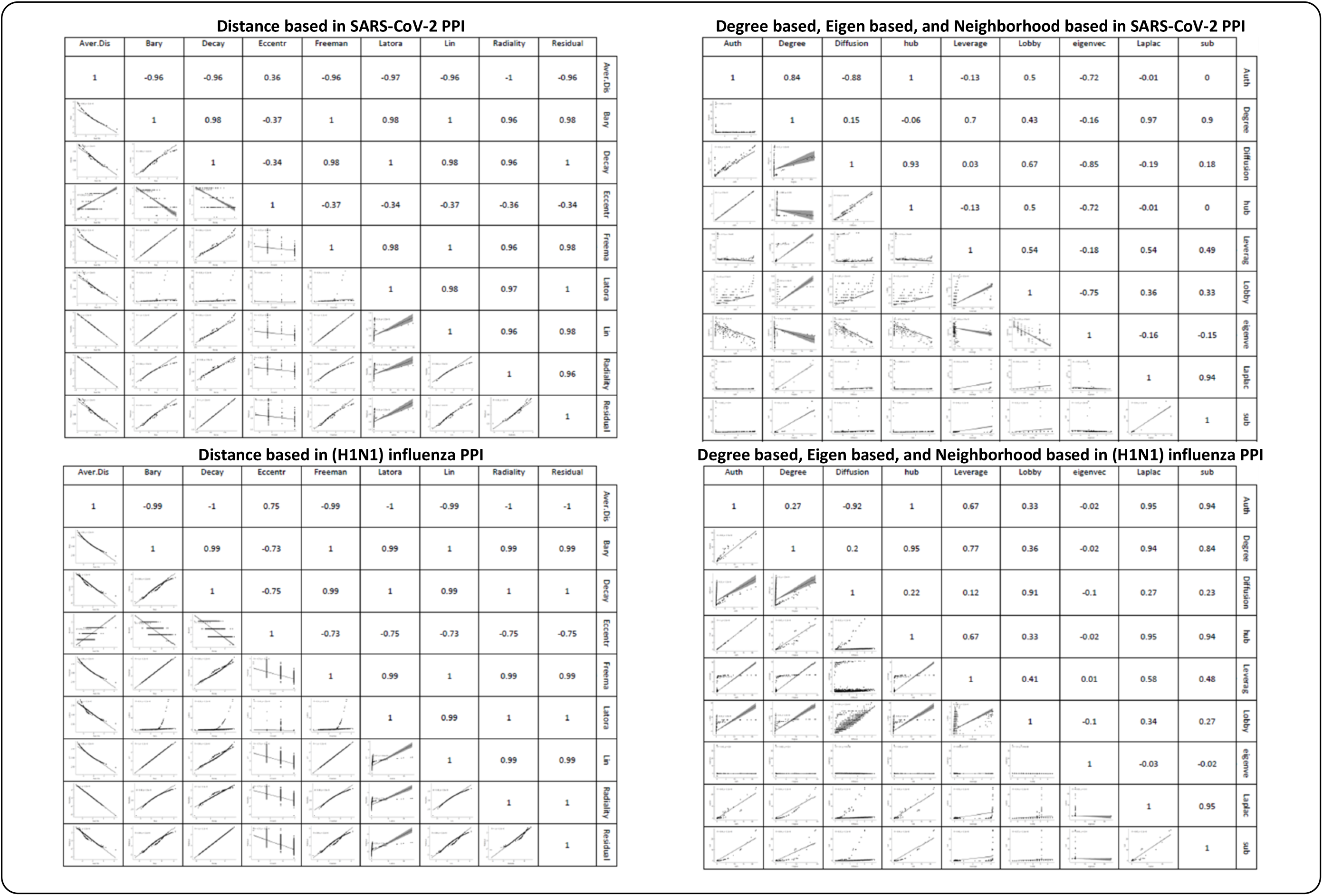
r Pearson correlation coefficients between centralities in two groups and pairwise scatter plots of centrality measures.

**Fig. 3.**
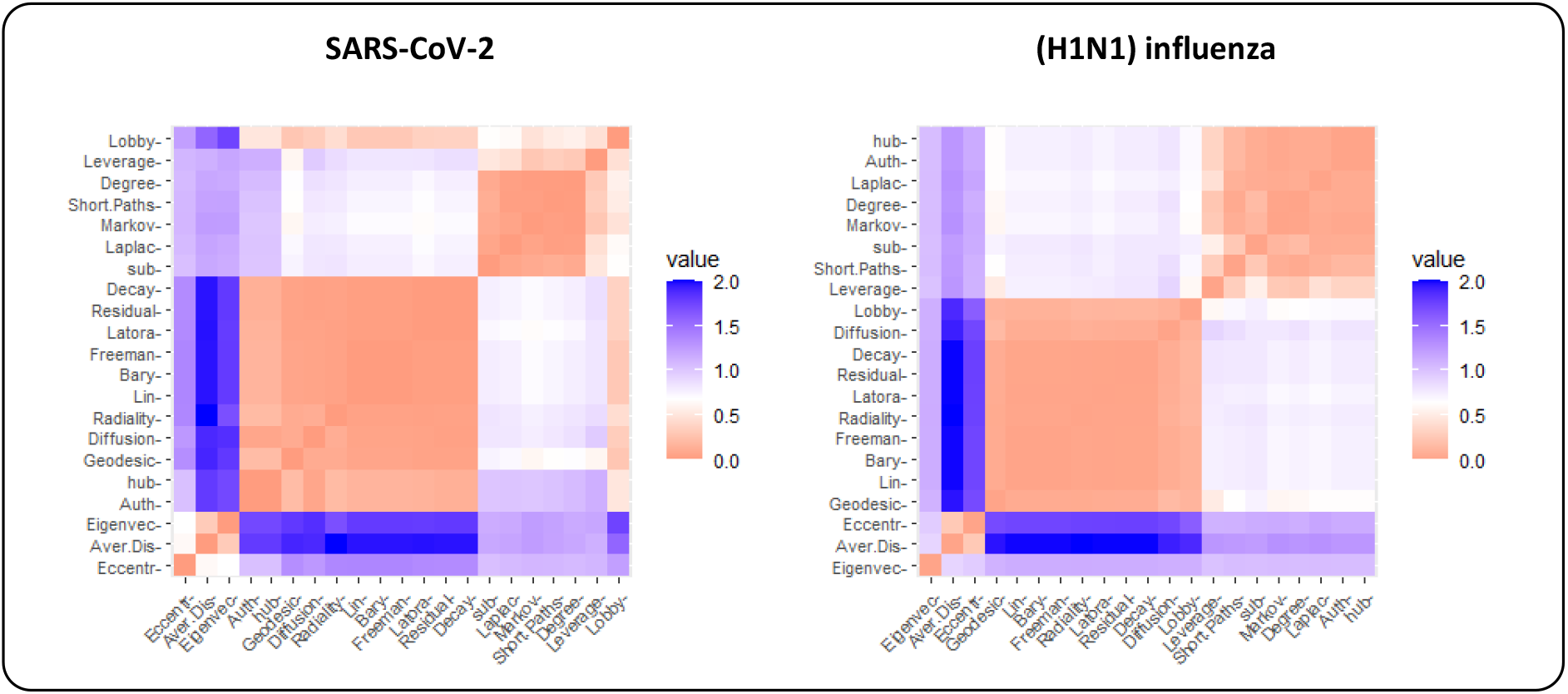
The dissimilarity matrix based on the Pearson correlation coefficient for all centrality measures in both networks

### Dimensionality reduction and clustering analysis

In the next phase, PCA-based dimensionality reduction was applied on centrality measures to show a visual representation of the dominant centrality measures in the data set. The profile of the distance to the center of the plots and their directions were mostly harmonic for both networks as illustrated in Fig.4. The contribution of each centrality measure for two dimensions is shown in Additional file 1. The percentage of contribution of variables (i.e. centrality measures) in a given PC was computed as (variable. Cos2*100)/(total Cos2 of the component)). Fig.5 illustrates the first ten contributing centrality measures to PCA for two dimensions. In both networks, the contribution percent for the first ten contributors is too close for the first dimension. For the second dimension, degree centrality is the major contributor for both PPINs. Eigenvector and Eccentricity revealed a low contribution value in both PPINs. In contrast, Closeness (Latora) displayed high levels of contribution in both networks whilst it was the first rank of SARS-CoV-2 PPIN contributors and second rank of (H1N1) influenza PPIN contributors.

**Fig. 4.**
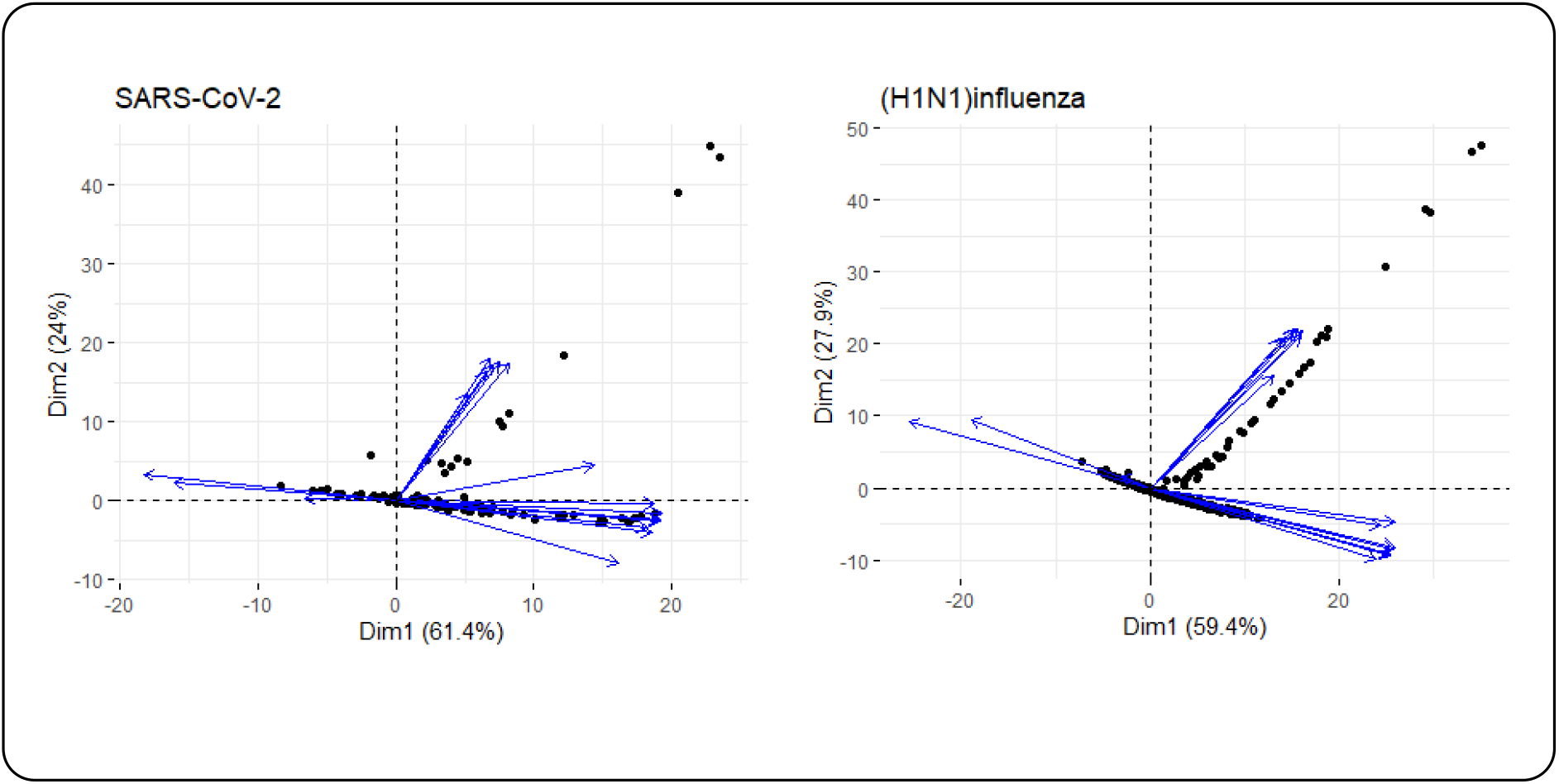
Biplot representation of the centrality measures in SARS-CoV-2 and (H1N1) influenza PPINs. In each plot, nodes were shown as points and centrality measures as vectors

**Fig.5.**
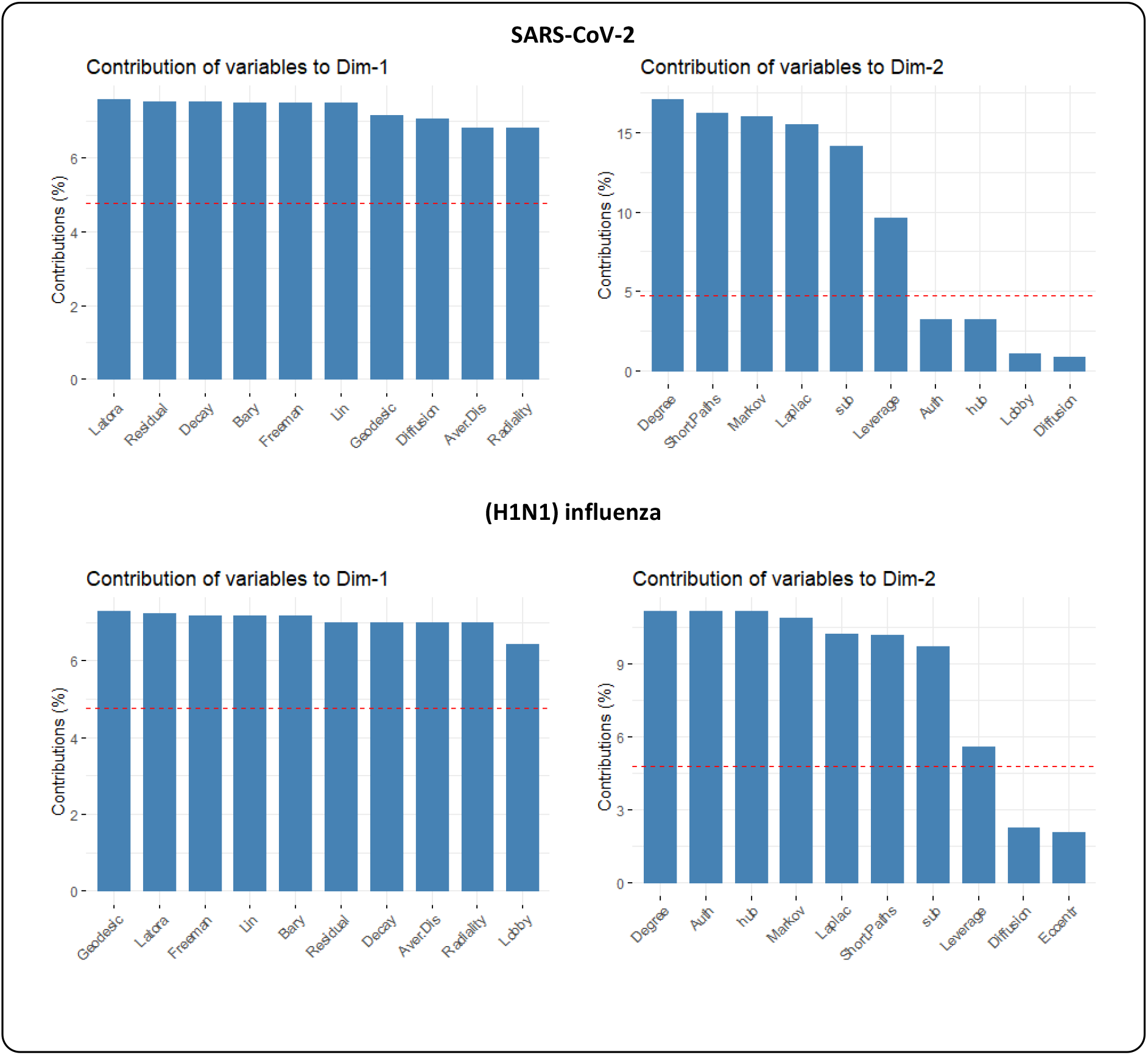
10 top contributing centrality measures to PCA for two dimensions

Ultimately, we performed unsupervised classification to cluster centrality values computed in PPINs. First, we executed a clustering tendency procedure. For clustering centrality values in each network, we considered Hopkins statistics were more than the threshold. The threshold value was 0.05 [11]. The results are shown in the first column of Table 5 and Additional file 2. Then, silhouette scores were calculated in three methods (i.e. hierarchical, k-means, and PAM) and average Silhouette width were evaluated in clustering the data sets (Additional file 3). Finally, based on average Silhouette width, the k-means method was selected for clustering centrality values in both PPINs (Fig.6). The outputs of clustering method and the corresponding number of clusters were also shown in Table 5. Optimal number of clusters were also determined by k-means and PAM clustering algorithms (Additional file 4). The centrality measures were clustered in each PPINs using the hierarchical algorithm based on Ward’s method [45] that were shown in Fig.7.

**Table 5.** Clustering information values for PPINs.

| Network | Hopkins Statistic | Number of Clusters | Average Silhouette width |
| --- | --- | --- | --- |
| SARS-CoV-2 | 0.75 | 9 | 0.42 |
| (H1N1) influenza | 0.77 | 10 | 0.36 |

**Fig.6.**
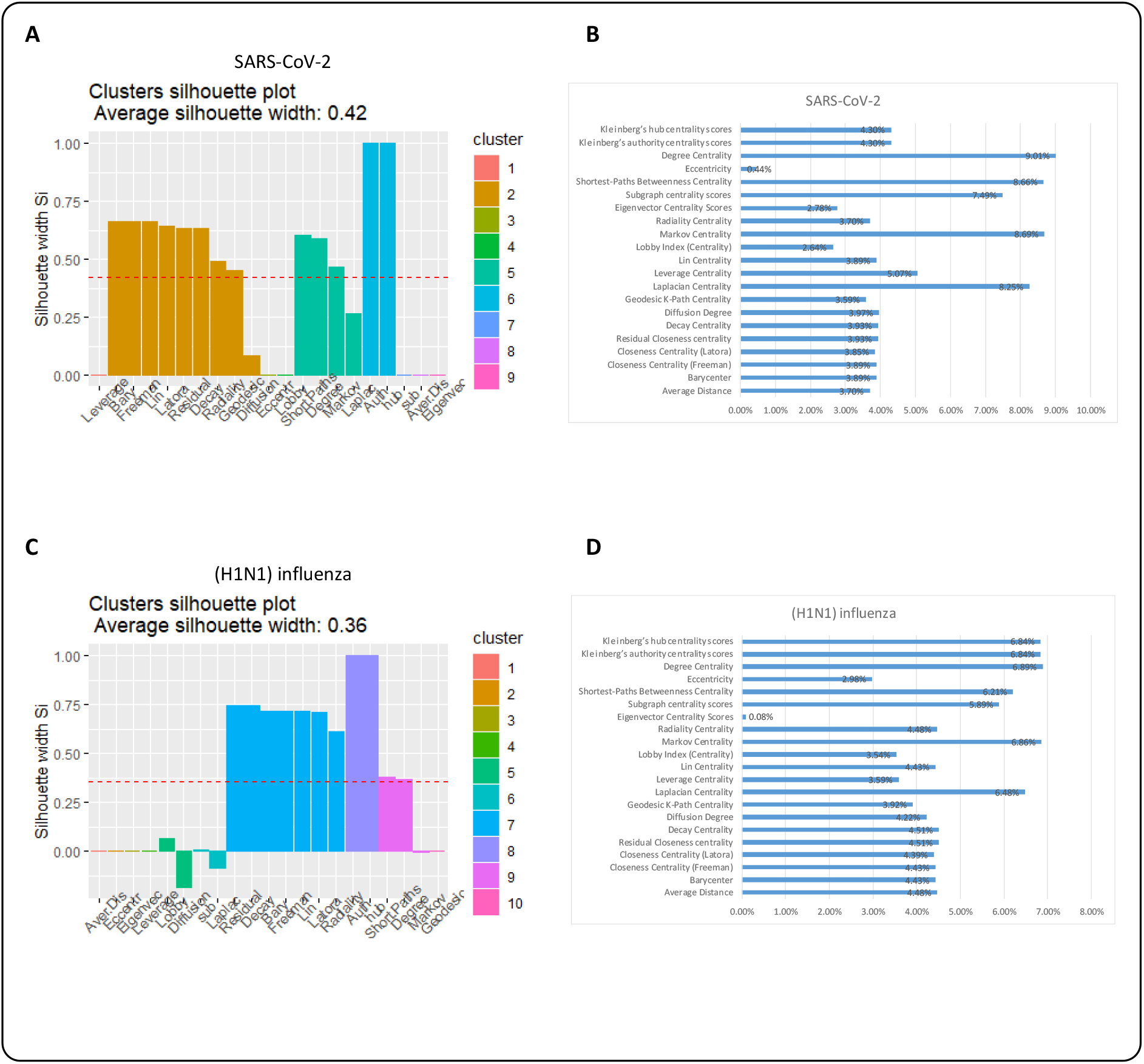
(A) Clustering silhouette plot of the combined-score PPIN. The colors represented the nine clusters of the centrality measures in SARS-CoV-2 PPIN. The average silhouette width was 0.42. (B) Contribution values of centrality measures according to their corresponding principal components in SARS-CoV-2 PPIN. (C) Clustering silhouette plot of the combined-score PPIN. The colors represented the ten clusters of the centrality measures in (H1N1) influenza. The average silhouette width was 0.36. (D) Contribution values of centrality measures according to their corresponding principal components in (H1N1) influenza PPIN.

**Fig. 7.**
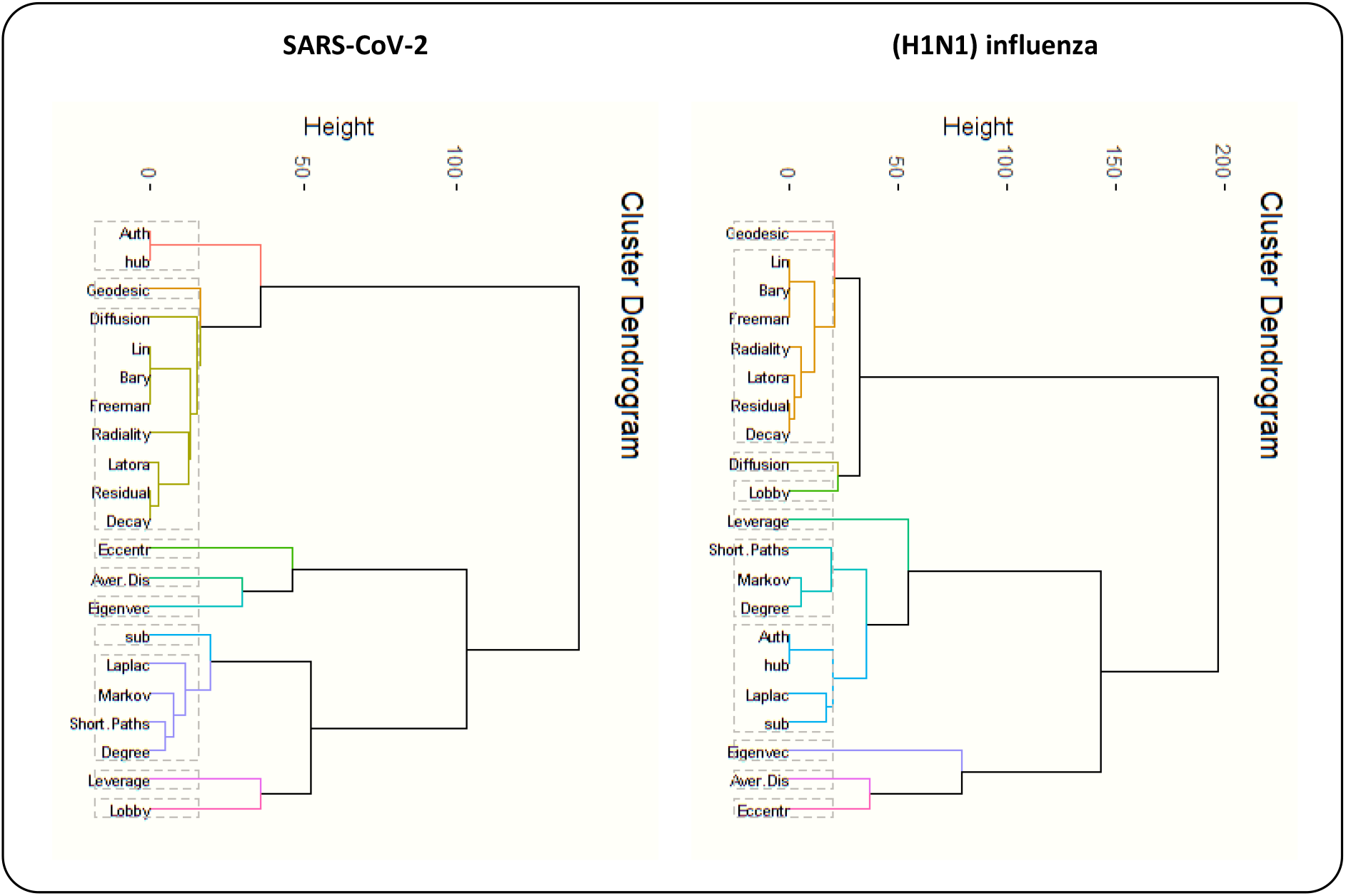
Clustering dendrograms for SARS-CoV-2 and (H1N1) influenza PPINs

## Discussion

At the validation step, we encountered remarkable results. Silhouette scores of centrality measures illustrated the centrality measures in the same clusters had very close contribution values for these measures (Fig. 6). In SARS-CoV-2 PPIN, Barycenter, Decay, Diffusion degree, Closeness (Freeman), Geodesic K-Path, Closeness (Latora), Lin, Radiality, and Residual closeness measures were in the same cluster. Also, in (H1N1) influenza, Barycenter, Decay, Closeness (Freeman), Closeness (Latora), Lin, Radiality, and Residual closeness were measures were in the same cluster. The average silhouette scores were 0.55 and 0.71 in these clusters for SARS-CoV-2 and (H1N1) influenza PPINs, respectively. The centrality measures namely Shortest-Paths betweenness, Laplacian, Degree, and Markov measures were in a cluster for SARS-CoV-2 PPIN where the mean of their silhouette scores (i.e. 0.48) was higher than the overall average, and in the same way, their corresponding contribution values were high, too. Kleinberg’s hub and Kleinberg’s authority scores are grouped in a cluster in both PPINs and their corresponding contribution values were equal.

Our results demonstrated that an exclusive profile of centrality measures including Barycenter, Decay, Diffusion degree, Closeness (Freeman), Closeness (Latora), Lin, Radiality, and Residual closeness was the most significant index to determine essential nodes. We inferred that both PPINs have close results in centrality analysis. Also, our research confirmed an analogous study [11] about the relationship between contribution value derived from PCA and silhouette width as a cluster validation. Furthermore, our centrality analysis resulted in many equal values in all centrality measures, this confirms dynamic robustness for PPI networks. Also, it reveals that PPI networks due to sparsity and tree-like topology are more explorable than random networks with higher connectivity [46].

## Conclusions

SARS-CoV-2, a novel coronavirus mostly known as COVID-19, has become a matter of critical concern around the world. Besides, network-based methods have emerged to analyze, and understand complex behavior in biological systems with a focus on topological features. In recent decades, network-based ranking methods have provided systematic analysis for predicting influence proteins and proposing drug target candidates in the treatment of types of cancer and biomarker discovery. SARS-CoV-2 and (H1N1) influenza PPI networks have 553 common proteins. Studying and comparing these networks can be an effective step to help biologists in drug design.

In this study, we studied SARS-CoV-2 and (H1N1) influenza PPI networks. Subsequently, 21 centrality measures were used for the prioritization of the proteins in both networks. We illustrated that dimensionality reduction methods like PCA can help to extract more relevant features (i.e. centrality measures) and corresponding relationships in unsupervised machine learning methods. Thus, to detect influential nodes in biological networks, PCA can help to select suitable measures. In other word, Dimensionality reduction methods can illuminate which measures have the highest contribution values, i.e., which measures contain much more useful information about centrality.

## Acknowledgement

We would like to acknowledge the help that received from our colleagues in Machine Learning and Bioinformatics Laboratory (MLBL) of University of Zanjan, Zanjan, Iran.

